# Transcriptome analysis indicates dominant effects on ribosome and mitochondrial function of a premature termination codon mutation in the zebrafish gene *psen2*

**DOI:** 10.1101/2020.04.20.050815

**Authors:** Haowei Jiang, Stephen Martin Pederson, Morgan Newman, Yang Dong, Michael Lardelli

## Abstract

*PRESENILIN 2* (*PSEN2*) is one of the genes mutated in early onset familial Alzheimer’s disease (EOfAD). *PSEN2* shares significant amino acid sequence identity with another EOfAD-related gene *PRESENILIN 1* (*PSEN1*), and partial functional redundancy is seen between these two genes. However, the complete range of functions of *PSEN1* and *PSEN2* is not yet understood. In this study, we performed targeted mutagenesis of the zebrafish *psen2* gene to generate a premature termination codon close downstream of the translation start with the intention of creating a null mutation. Homozygotes for this mutation, *psen2*^*S4Ter*^, are viable and fertile, and adults do not show any gross pigmentation defects, arguing against significant loss of γ-secretase activity. Also, assessment of the numbers of Dorsal Longitudinal Ascending (DoLA) interneurons that are responsive to *psen2* but not *psen1* activity during embryogenesis did not reveal decreased *psen2* function. Transcripts containing the *S4Ter* mutation show no evidence of destabilization by nonsense-mediated decay. Forced expression in zebrafish embryos of fusions of *psen2*^*S4Ter*^ 5’ mRNA sequences with sequence encoding enhanced green fluorescent protein (EGFP) indicated that the *psen2*^*S4Ter*^ mutation permits utilization of cryptic, novel downstream translation start codons. These likely initiate translation of N-terminally truncated Psen2 proteins that obey the “reading frame preservation rule” of *PRESENILIN* EOfAD mutations. Transcriptome analysis of entire brains from a 6-month-old family of wild type, heterozygous and homozygous *psen2*^*S4Ter*^ female siblings revealed profoundly dominant effects on gene expression likely indicating changes in ribosomal, mitochondrial, and anion transport functions.

## Introduction

*PRESENILIN 2* (*PSEN2*) was first identified as a candidate locus for mutations causing familial Alzheimer’s disease (AD) with early onset (EOfAD) when a point mutation resulting in the substitution of an isoleucine residue for an asparagine residue (N141I) was found in a Volga German AD family in 1995 (1). *PSEN2* is similar in structure to the major EOfAD gene *PRESENILIN 1* (*PSEN1*) and the two genes encode proteins with 62% amino acid sequence identity (2). The age of onset of Alzheimer’s disease (AD) caused by mutations in *PSEN2* ranges from 39 to 75 years, which overlaps both with *PSEN1* EOfAD-associated mutation disease onset ages and with late onset, sporadic AD (3). The later mean onset age of AD caused by *PSEN2* mutations compared to mutations in *PSEN1* is still unexplained, but some studies suggest that it may be caused by the partial replacement of *PSEN2* function by *PSEN1* (4). However, the functions of *PSEN1* and *PSEN2* have not yet been determined comprehensively. Moreover, despite the partial functional redundancy between *PSEN1* and *PSEN2, in vitro* studies have shown that the protein products of the two genes also play divergent roles in cellular physiology ((5, 6) and reviewed in (7)).

Both PSEN1 and PSEN2 proteins are components of γ-secretase complexes. The absence of PSEN1 is thought to reduce γ-secretase activity in mammalian cells (8, 9), while the absence of both PSEN1 and PSEN2 is thought to eliminate it completely (10, 11) although some data does not agree with this (reviewed in Jayne et al. (12)). In mice, the loss of Psen1 causes premature differentiation of neural progenitor cells (NPC) and inhibition of Notch signaling leading to skeletal defects (13), and, ultimately, perinatal lethality (14). Mouse embryos lacking both Psen1 and Psen2 activity are more severely affected, showing earlier lethality and a developmental phenotype similar to loss of Notch1 activity (15, 16). However, by itself, the absence of Psen2 activity in mice does not appear to affect development significantly (17). In zebrafish, the inhibition of either Psen1 or Psen2 translation caused decreased melanocyte numbers in trunk and tail and other effects of decreased Notch signaling indicating a possibly greater role in Notch signaling for Psen2 protein in zebrafish compared to in mammals (18). Inhibition of Psen2 translation also led to increased Dorsal Longitudinal Ascending (DoLA) interneuron number, while inhibition of Psen1 translation showed no effect on this neuronal cell type (18).

Although mammalian *Psen1* and *Psen2* show compensatory regulation with forced down-regulation of one causing up-regulation of the other (19, 20), only *Psen2* down-regulation causes markedly decreased γ-secretase activity in the microglial cells of mice. The inhibition of γ-secretase activity caused by forced down-regulation of *Psen2* led to exaggerated proinflammatory cytokine release from microglia, indicating that *Psen2* plays an important role in central nervous system innate immunity (20). Furthermore, a negative regulator of monocyte pro-inflammatory response, miR146, was found to be constitutively down-regulated in the microglia of a *Psen2* knockout mouse strain, supporting that *Psen2* dysfunction may be involved in neurodegeneration through its impacts on the pro-inflammatory behavior of microglia (21). Also, *Psen2* (but not *Psen1*) knockout mice show reduced responsiveness to lipopolysaccharide as well as decreased expression of nuclear factor kappa-light-chain-enhancer of activated B cells (NF-kappaB), reduced mitogen-activated protein kinase (MAPK) activity and reduced pro-inflammatory cytokine production. This indicates that *Psen2* has a specific function(s) in innate immunity independent of *Psen1* (22).

The particular role of mammalian Psen2 protein in inflammation is consistent with its restricted localization to the mitochondrial associated membranes (MAM) of the endoplasmic reticulum (ER) (23). MAM formation has been shown to influence inflammatory responses, is the site of autophagosome initiation, and plays a major role in regulating mitochondrial activity (reviewed in Marchi et al (24)).

Considerable evidence supports roles for PRESENILIN proteins in the function of mitochondria (25), which are central to energy production in cells and to other cellular processes affected in AD such as apoptosis, reactive oxygen species production, and calcium homeostasis (26). In human cell lines, it has been reported that PSEN2, but not PSEN1, modulates the shuttling of Ca^2+^ between the ER and mitochondria since mitochondrial Ca^2+^ dynamics are reduced by PSEN2 down-regulation and enhanced by the expression of mutant forms of PSEN2 (27). In mouse cell lines, deficiency of *Psen2* led to reduced expression of subunits responsible for mitochondrial oxidative phosphorylation with altered morphology of the mitochondrial cristae, as well as an increase in glycolytic flux. This indicated that absence of Psen2 protein causes an impairment in respiratory capacity with a corresponding increase in glycolytic flux to support cells’ energy needs (28).

Despite the identification of hundreds of different EOfAD mutations in human *PSEN1* and *PSEN2*, none of these appear to remove all gene function (i.e. none are null mutations)(12). All *PRESENILIN* EOfAD mutations follow the “reading frame preservation rule” meaning that they all produce at least one transcript variant containing an open reading frame (ORF) termined by the original (non-mutant) stop codon (12).

As part of an effort using zebrafish to identify the specific cellular changes caused by EOfAD-like mutations in these genes, we wished to examine null mutations so that their effects could be excluded from consideration. In this paper we describe an unsuccessful attempt to generate a null mutation of the zebrafish orthologue of the human *PSEN2* gene, *psen2*, by introduction of a premature termination codon downstream of the asssumed translation start codon. Unexpectedly, the mutation appears to force utilization of downsteam methionine codons for translation initiation. This generates N-terminally truncated proteins that act dominantly in an EOfAD-like manner.

## Materials and Methods

### Animal ethics

All experiments using zebrafish were conducted under the auspices of the Animal Ethics Committee of the University of Adelaide. Permits S-2014-108 and S-2017-073.

### sgRNA design and synthesis

The target sequence of Ps2Ex3 sgRNA is 5’-CAGACAGTGAAGAGGAC *TCC*-3’. This target sequence was cloned into the plasmid pDR274 (Addgene plasmid # 42250) (29). The Ps2Ex3 pDR274 plasmid was linearized with *HindIII-HF* ^®^ (NEB, Ipswich, Massachusetts, USA, R3104S), and then used as a template for synthesis of Ps2Ex3 sgRNA with the MAXIscript™ T7 Transcription Kit (Ambion, Inc, Foster City, California, USA, AM1312).

### Injection of zebrafish embryos

Tübingen (wild type embryos were generated by mass mating. Ps2Ex3 sgRNA (90 ng/μL final concentration) was first mixed with Cas9 nuclease (Invitrogen, Carlsbad, California, USA, B25640), and then incubated at 37°C for 15 min to maximize cleavage efficiency. 5-10 nL of the mixture was injected into zebrafish embryos at the one-cell stage. The injected embryos were subsequently raised for mutation screening.

### Mutation detection in G0 injected embryos using T7 Endonuclease I

Mutation detection was based on the T7 Endonuclease I assay (30). Since mismatches, small insertions or deletions generated through non-homologous end joining (NHEJ) result in failure of base-pairing in heteroduplexes at mutation sites, T7 Endonuclease I is able to recognize and cleave at the sites of these mutations.

To test whether the CRISPR/Cas9 system had functioned in the injected G0 embryos, 10 embryos were randomly selected from each injected batch and pooled together for genomic DNA extraction at ∼24 hours post fertilization (hpf). To extract the genomic DNA, these 10 embryos were placed in 100 μL of 50 mM NaOH and then heated to 95°C for 15 min, and then 1/10th volume of 1 M Tris-HCl, pH 8.0 was added to each sample to neutralize the basic solution after cooling to 4°C (31). A pair of primers (5’-AGGCCACATCACGATACAC -3’ and 5’-TGACCCGTTTGCTGTCTG-3’) binding to the flanking regions of the intended cleavage site was designed to amplify the test region (∼472 bp) through PCR. The PCR conditions for this amplification reaction were 95°C, 2 min; 31 cycles of [95°C, 30 s; 58°C, 30 s; 72°C, 30 s]; then 72°C, 5 min. The PCR products were purified using the Wizard® SV Gel and PCR Clean-Up System (Promega, Wisconsin, USA, A9281) and annealed (denaturation at 95°C for 5 min and then slow cooling of the samples at the rate of -2°C/sec from 95°C to 85°C and then - 0.1°C/sec from 85°C to 25°C for annealing of heteroduplexes) before addition of T7 Endonuclease I (NEB, Ipswich, Massachusetts, USA, M0302S). Heteroduplexes containing small mutations at the intended site should be cleaved into two fragments, ∼313 bp (upstream) and ∼159 bp (downstream).

When the T7 Endonuclease I assay on injected G0 embryos showed the presence of mutation at the target site, the remaining embryos from the same injection batch were raised for further mutation screening and breeding.

### Mutation detection in adult G0 and F1 fish using T7 Endonuclease I and Sanger sequencing

When a G0 injected fish had grown to sufficient size (>2 cm in length, 2 to 3 months old), the tip of its tail (∼2 mm in length) was biopsied (clipped) under Tricaine (1.68μg/mL) anesthesia for genomic DNA extraction. The clipped tail was placed in 100 μL of 50 mM NaOH and then heated to 95°C for 15 min to extract genomic DNA. The sample was then cooled to 4°C, and a 1/10th volume of 1 M Tris-HCl, pH 8.0 was then added to each sample to neutralize the basic solution (31). The same T7 Endonuclease I assay used previously for mutation detection in G0 embryos was then applied to the genomic DNA extracted from the G0 adult fish biopsy. However, since each G0 mutation-carrying fish was probably mosaic for several different mutations at the target site, each G0 fish was outbred to a wild type Tübingen fish, to produce the F1 progeny, some of which could be heterozygous for single mutations. The F1 fish were biopsied and screened using the T7 Endonuclease I assay when large enough. For F1 fish found to carry mutations, the PCR-amplified fragments were sent to the Australian Genome Research Facility (AGRF, North Melbourne, VIC, Australia) for **S**anger sequencing to identify the mutations.

An 8-bp deletion resulting in a frameshift downstream of the start codon of *psen2, psen2*^*S4Ter*^ (Figure 1), was identified. PCR primers specifically detecting this mutation were designed (*psen2*^*S4Ter*^ forward primer: 5’-TTCATGAATACCTGAAGAGG-3’, wild type forward primer: 5’-TTCATGAATACCTCAGACAGTG-3’, and reverse primer: 5’-GAACAGAGAATGTACTGGCAGC-3’) for further screening. The PCR conditions for *psen2*^*S4Ter*^ mutant detection are 95°C, 2 min; 31 cycles of [95°C, 30 s; 55°C, 30 s; and then 72°C, 30 s]; 72°C, 5 min. The length of PCR products is ∼230 bp. The PCR conditions for wild type-specific detection are 95°C, 2 min; 31 cycles of [95°C, 30 s; 60°C, 30 s; and 72°C 30 s]; 72°C, 5 min and the anticipated length of the PCR products is ∼230 bp.

**Figure 1.**
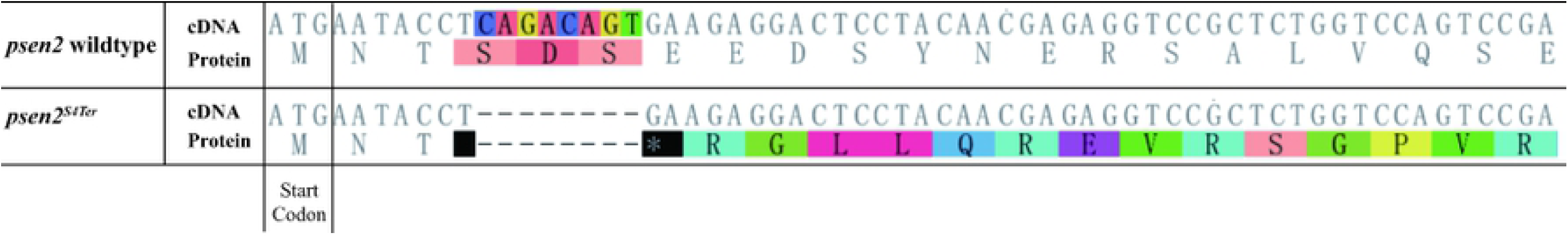
Predicted protein sequence of *psen2*^*S4Ter*^. An 8-bp deletion resulted in a frameshift downstream of the nominal translation start codon of *psen2* creating a stop codon as the 4^th^ codon.

### Breeding of *psen2*^*S4Ter*^ mutant fish

The initial F1 fish carrying the *psen2*^*S4Ter*^ mutation was outbred to a wild type fish to generate a population of F2 progeny that was 50% heterozygous mutants and 50% wild type fish. Two F2 heterozygous mutant fish were then inbred to generate a family of F3 fish consisting of (theoretically) 50% heterozygous mutants, 25% homozygous mutants and 25% wild type fish. This F3 family was raised to six months of age before brain removal and total brain RNA extraction for RNA-seq and other analyses.

### Total RNA extraction from 6-month-old zebrafish brains

Individual fish were genotyped using PCR. Fish of the desired genotype were then selected and euthanized by sudden immersion in an ice water slurry for at least 30 seconds before immediate decapitation. The entire brain was then removed from the cranium for extraction of total RNA for either digital quantitative PCR (dqPCR) on cDNA or RNA-seq (below).

For dqPCR tests, six wild type, six heterozygous and six homozygous fish from the F3 family were selected. Three of each genotype were then exposed to acute hypoxia (dissolved oxygen content of the water was ∼1.0 mg/L) for ∼2.5 h, while the remaining three of each genotype were exposed to normoxia. Fish were then euthanized and brains removed (as above) for total RNA was extracted using the RNeasy Mini Kit (QIAGEN, Venlo, Netherlands, 74104). cDNA was synthesised from brain RNAs using the SuperScript™ III First-Strand Synthesis System (Invitrogen, Carlsbad, California, USA, 18080051) with Random Primers (Promega, Madison, Wisconsin, USA, C1181).

For RNA-seq, four wild type, four *psen2*^*S4Ter*^ heterozygous and four *psen2*^*S4Ter*^ homozygous mutant brains (all from female fish) were extracted from the same family. Total RNA from these brains was extracted using the *mir*Vana™ miRNA Isolation Kit (Ambion, Inc, Foster City, California, USA, AM1560). RNA samples were sent to the Australian Cancer Research Foundation (ACRF) Cancer Genomics Facility, Adelaide SA, Australia for sequencing.

### Allele specific transcript expression analysis by dqPCR

Primers for dqPCR, including a reverse primer specifically detecting the wild type allele (5’-TCGTTGTAGGAGTCCTCTTCACTG-3’), a reverse primer specifically detecting the *psen2*^*S4Ter*^ allele (5’-TCGTTGTAGGAGTCCTCTTCAGG-3’) and a common forward primer (5’-TTCCTCACTGAATTGGCGATG-3’), were designed for allele specific expression analysis of the F3 family using the QuantStudio™ 3D Digital PCR System (Life Sciences, Waltham, MA, USA) with the QuantStudio™ 3D Digital PCR 20K Chip Kit v2 and Master Mix (Life Sciences, Waltham, MA, USA, A26317) and SYBR™ Green I Nucleic Acid Gel Stain (Life Sciences, Waltham, MA, USA, S7563). The dqPCR conditions for assays of mutant allele or wild type allele expression were 96°C, 10 min; 49 cycles of [62°C, 2 min; 98°C, 30 s]; 62°C, 2 min. The lengths of the anticipated PCR products are ∼130 bp. 25 ng of total cDNA^*^ from a sample was loaded into one chip for the dqPCR. The chips were read using QuantStudio™ 3D AnalysisSuite Cloud Software (Life Sciences, Waltham, MA, USA).

^*^Stated cDNA concentrations are based on measured concentrations of RNA under the assumption that subsequent reverse transcription is completely efficient.

### Testing for aberrant *psen2* transcript splicing due to the *S4Ter* mutation

Fifteen 24-hour-old zebrafish embryos were collected into one tube. Total RNA was extracted using the QIAGEN RNeasy mini Kit (QIAGEN, Hilden, Germany). 400ng of total RNA from each brain was then used to synthesize 20μL of first-strand cDNA by reverse transcription (SuperScript III kit, Invitrogen, Camarillo, California, USA). 40ng of each cDNA preparation (a quantity calculated from the RNA concentration on the assumption that reverse transcription of RNA into cDNA was complete) was used to perform PCR using GoTaq® DNA polymerase (Promega, Madison, USA). Each 25μL PCR reaction contained 0.2mM of deoxyribonucleotide triphosphates (dNTPs), 0.4μM of each PCR primer, 1 unit of GoTaq® DNA polymerase and 40ng of zebrafish embryo cDNA template. PCR cycling was performed with 35 cycles of a denaturation temperature of 95°C for 30s, then an annealing temperature of 60°C for 45s and then an extension temperature of 72°C for 1.5 minutes. PCR products were electrophoresed through a 1.5% agarose gel in 1×TAE buffer for separation and identification. **PCR primers:** 5’UTR_F: 5’-TTTTGACGGAGTATTTTCGCAT-3’, Exon3_F: 5’-CTCTTCATCCCTGTCACGCTCT-3’, Exon3_R: 5’-CTCGGTGTAGAAACTGACGGACTT-3’, Exon7_R: 5’-TTCCACCAGCATCCTCAACG-3’.

### Construction of the *psen2* EGFP fusion expression vectors

DNA sequence corresponding to the 5’UTR and the first 113 codons of the *psen2* gene fused to sequence encoding the N-terminal end of EGFP (but excluding EGFP’s translation start codon) was synthesized by Integrated DNA Technologies Inc. Coralville, Iowa, USA) and ligated into the pcGlobin2 vector between the *BamH* I and *EcoR* I restriction sites to construct expression vector psen2WT-EGFP. The same procedure was followed to construct expression vector psen2S4Ter-EGFP that is identical to psen2WT-EGFP except that psen2S4Ter-EGFP lacks the 8 nucleotides deleted in the *psen2*^*S4Ter*^ mutant allele (i.e. 5’-CAGACAGT -3’). The complete sequences of these fusion genes are given in supplementary data file S1 Appendix 1.

### *In-vitro* mRNA transcription and microinjection

The psen2WT-EGFP and psen2S4Ter-EGFP expression vector constructs were restricted with *Xba*I before transcription using the mMessage mMachine T7 kit (Thermo Fisher Scientific Inc, Waltham, Massachusetts, USA) to generate mRNA. All mRNAs were precipitated with LiCl and then redissolved in water for injection of 2-5 nL at a concentration of 400 ng/μL. No obvious developmental abnormalities were seen after injection of these mRNAs into zebrafish embryos. At ∼24 hpf, both the psen2WT-EGFP mRNA- and the psen2S4Ter-EGFP mRNA-injected embryos showed weak EGFP fluorescence (as visualized by fluorescence microscopy). For mRNA injected and non-injected embryos (as negative controls) 15 embryos were collected for subsequent western immunoblot analysis.

### Western immunoblot analyses

Embryos at 24 hpf were first dechorionated and their yolks were removed in embryo medium containing Tricaine methanesulfonate. Embryos were then placed in RIPA extraction buffer (Sigma-Aldrich Corp. St. Louis, Missouri, USA) containing Complete Proteinase Inhibitor (Sigma-Aldrich), homogenized and incubated at 4°C (with rotation). Cell debris was sedimented from the protein sample by centrifugation at 16,000 x g for 30 seconds, LDS sample buffer (Invitrogen) was added to the supernatant, the protein sample was heated at 80°C for 20 minutes and then stored at -80°C. Sample Reducing Agent (Thermo Fisher Scientific) was added to samples prior to being loaded onto NuPAGE™ 4-12% Bis-Tris Protein Gels (Invitrogen). The separated proteins were subsequently transferred to a PVDF membrane (Bio-Rad Laboratories, Hercules, California, USA) using the Mini Gel Tank and Blot Module Set (Thermo Fisher Scientific). For EGFP protein detection, the PVDF membrane was blocked with blocking reagent (Roche Holding AG, Basel, Switzerland) and then probed with the primary antibody, polyclonal anti-GFP goat (Rockland Immunochemicals Inc., Gilbertsville, Pennsylvania, USA), followed by secondary antibody, horseradish peroxidase (HRP) conjugated anti-goat antibody (Rockland). For subsequent beta-tubulin (protein loading control) detection, the PVDF membrane was blocked in 5% skim milk and then probed with the primary antibody, monoclonal beta-tubulin (E7, DSHB) followed by secondary antibody, horseradish peroxidase conjugated anti-mouse antibody (Rockland). Bound antibody was detected by chemiluminescence using SuperSignal™ West Pico PLUS Chemiluminescent Substrate (Thermo Fisher Scientific) and imaged by the ChemiDoc™ MP Imaging System (Bio-Rad Laboratories).

### RNA-seq data generation and quality control

Paired-end (2×150bp) RNA Seq libraries were provided by the ACRF Cancer Genomics Facility. Briefly, RNA samples were depleted for rRNA using the methods of Adiconis (32) and sequences derived from mammalian rRNA, before being prepared using the KAPA Hyper RNA Library Prep kit (Roche Holding AG), and sequenced on an Illumina NextSeq 500 (Illumina, San Diego, California, USA). Libraries were generated for *n = 4* samples from each of the genotypes: wild type (WT, *psen2*^*+/+*^), heterozygous (Het, *psen2*^*S4Ter*^*/+*) and homozygous (Hom, *psen2*^*S4Ter*^*/psen2*^*S4Ter*^), ranging in size from 27,979,654 to 37,144,975 reads. Libraries were trimmed using cutadapt v1.14 to remove Illumina Adapter sequences. Bases with a PHRED score < 30 were also removed along with NextSeq-induced polyG runs. Reads shorter than 35bp after trimming were discarded. Trimmed reads were aligned to GRCz11 using STAR v2.7.0 (33), and gene descriptions based on Ensembl release 98. For the purposes of genotype confirmation, trimmed reads were additionally aligned using kallisto v0.43.1 (34) to a modified version of the Ensembl transcriptome, where the sequence for the psen2S4Ter allele was additionally included. RNA-seq data has been deposited in the Gene Expression Omnibus database (GEO) under Accession Number GSE148468.

### Differentially expressed gene analysis of RNA-seq data

Unique alignments corresponding to strictly exonic regions from gene models in Ensembl release 98 were counted using featureCounts from the Subread package v1.5.2 (35), giving total counts per sample which ranged between 11,852,141 and 16,997,219. Genes with counts per million (CPM) > 1.5 in at least four samples were retained, whilst genes whose biotype corresponded to any type of rRNA were additionally excluded, giving 16,640 genes for differential expression analysis. As GC bias was suspected, given variable rRNA depletion across samples, gene-level counts were normalized for GC and length bias using CQN (36), before estimating fold change using the GLM likelihood-ratio test in the R package edgeR (37). Differential expression was determined for the presence of the mutant allele and the comparison between heterozygous and homozygous mutants. P-values from likelihood-ratio tests were adjusted using both the Benjamini-Hochberg FDR procedure, and the Bonferroni adjustment to provide two complementary view points on the data. Genes were considered to be differentially expressed in the presence of a mutant allele if satisfying one of two criteria, either 1) a Bonferroni-adjusted p-value < 0.01, or 2) an FDR-adjusted p-value < 0.01 along with a logFC estimate beyond the range ± 1. For the comparison between mutant genotypes, genes were considered to be exhibiting differential expression if obtaining an FDR-adjusted p-value < 0.05 in this comparison. Code for the complete analysis is available at https://uofabioinformaticshub.github.io/20170327_Psen2S4Ter_RNASeq/

### Genotype confirmation in RNA-seq data

Genotypes were confirmed for each RNASeq sample using transcript-level counts for the *psen2* wild-type allele and the *psen2*^*S4Ter*^ allele. No expression of the WT allele was overserved in any homozygous mutant, whilst no expression of the mutant allele was observed in homozygous WT samples. A ratio of approximately 1:1 between mutant and WT alleles was observed in all heterozygous mutant samples, as expected.

### Enrichment analysis

Enrichment testing was performed using Hallmark, KEGG and GO genesets from the MSigDB database (38), with GO terms excluded if the shortest path back to the root node was < 3 steps. For the comparison between mutant and wild-type samples, enrichment analysis was performed on the set of 615 DE genes using goseq (39), setting gene-level correlation with sample-specific rRNA content as a covariate. For the comparison between mutant genotypes, the number of DE genes was considered to be too small for this type of analysis. An additional enrichment analysis on the complete dataset was performed using fry on comparisons both between mutant and wild-type, and between mutant genotypes, as fry more appropriately handles inter-gene correlations than approaches like GSEA (40).

### *In situ* transcript hybridization analysis of DoLA neuron number

This was performed as previously described (41) on embryos from a pair mating of two *S4Ter* heterozygous mutants. After counting of DoLA neurons in an embryo, the embryo was subjected to DNA extraction as for the tail biopsies (above) and then its *psen2* genotype was determined by allele-specific PCRs.

## Results

### Generation of a putatively null mutation in zebrafish *psen2*

As part of a program analyzing the function of genes involved in familial Alzheimer’s disease, we wished to identify changes in the expression of genes in adult brains due to simple loss of *PSEN2* activity. We previously identified the *psen2* gene in zebrafish (42) and the ENSEMBL database (http://asia.ensembl.org) reports one *psen2* transcript (ENSDART00000006381.7) with 11 exons and the translation start codon residing in exon 2. Therefore, we used the CRISPR Cas9 system to generate a frameshift mutation just downstream of this transcript’s nominal translation start codon (intended to allow ribosomes to initiate translation but not translate Psen2 protein). A frameshift mutation (a deletion of 8 nucleotides) starting in the 4^th^ codon and resulting in the creation of a translation termination codon was isolated (Figure 1). The mutant allele is designated *psen2*^*S4Ter*^.

### No decreased stability of mutant allele transcripts under normoxia or hypoxia

Premature termination codons in transcripts frequently cause nonsense-mediated decay when more than 50-55 nucleotides upstream of an exon-exon boundary (43). Also, hypoxia appears to be an important element in AD (44, 45) and increases the expression of *PSEN* gene transcripts in both human and zebrafish cells (46, 47). Therefore, we sought to determine whether transcripts of the *psen2*^*S4Ter*^ allele are less stable than wild type transcripts and to observe the expression of mutant allele transcripts under acute hypoxia. We used dqPCR with allele-specific primer pairs to quantify relative transcript numbers in cDNA synthesized from the brains of 6-month-old wild type, heterozygous and homozygous mutant fish under normoxia or acute hypoxia (see Materials and Methods). The results of this analysis are shown in Figure 2. In heterozygous fish under normoxia, both wild type and mutant allele transcripts are expressed at similar levels in 6-month-old brains, with the wild type allele expressed at approximately half the level seen in wild type fish (i.e. that possess two wild type alleles). As expected, acute hypoxia increases the expression of the wild type transcript and this is also observed for the mutant transcript that shows no evidence of destablization (Figure 2).

**Figure 2.**
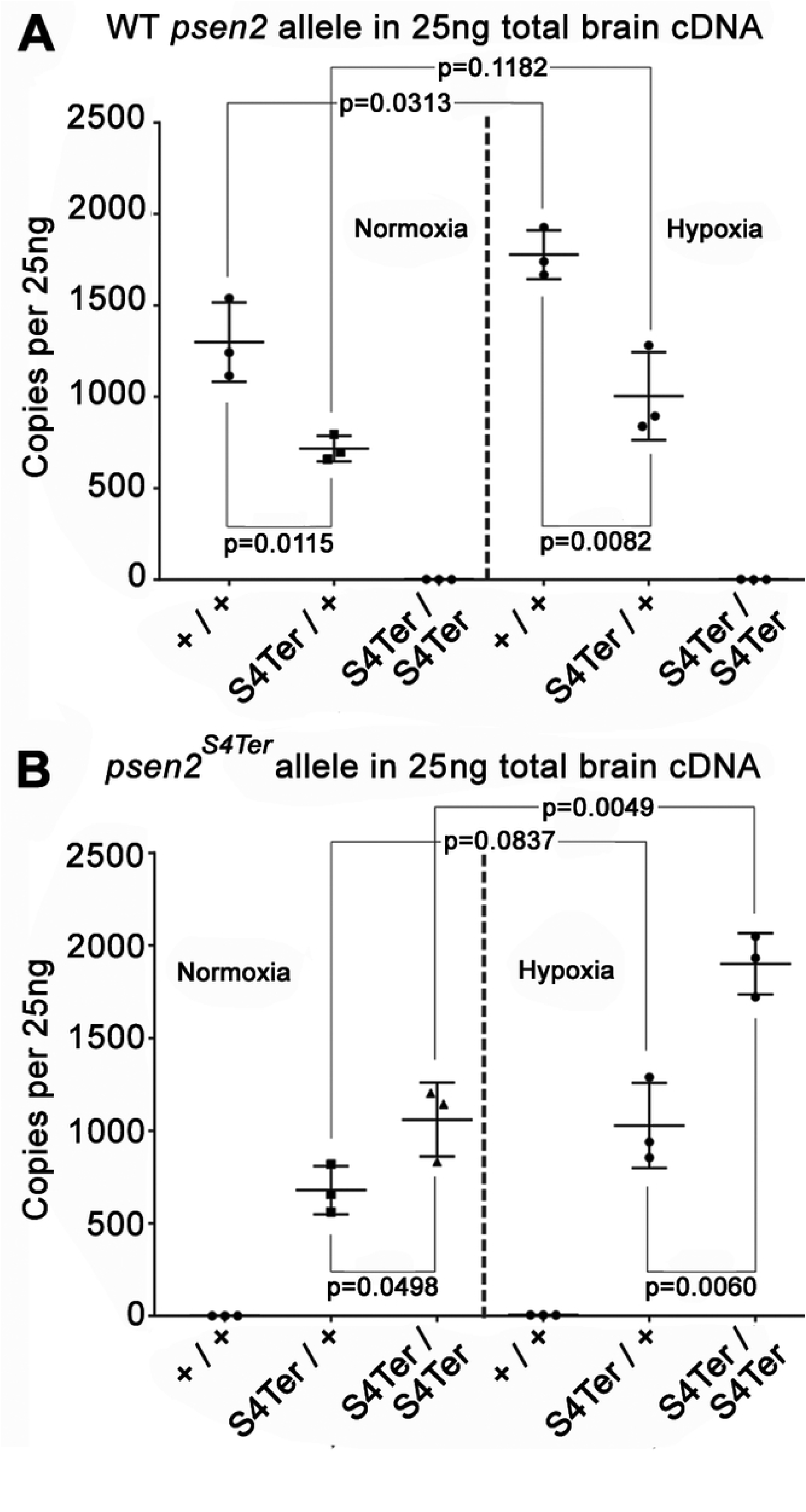
Allele-specific mRNA expression in the brains of 6-month-old fish of different genotypes under normoxia or acute hypoxia (as copies per 25 ng of brain cDNA in each digital qPCR) (A) The levels of wild type *psen2* allele mRNA in the *psen2*^*S4Ter*^/+ fish (∼700 copies) were significantly (p=0.0115) lower than in their wild type siblings (∼1,300 copies) under normoxia. Under hypoxia, the levels of wild type *psen2* allele mRNA in both the *psen2*^*S4Ter*^/+ fish (∼1,000 copies) and their wild type siblings (∼1,800 copies) were up-regulated, but only the higher levels in the wild type fish showed a statistically significant increase (p=0.0313) compared to the normoxic controls. (B) The levels of *psen2*^*S4Ter*^ allele mRNA in the *psen2*^*S4Ter*^/+ fish (∼700 copies) were significantly (p=0.0498) lower than in the *psen2*^*S4Ter*^/*psen2*^*S4Ter*^ fish (∼1,000 copies) under normoxia. Under hypoxia, the levels of *psen2*^*S4Ter*^ allele mRNA in both the *psen2*^*S4Ter*^/+ fish (∼1,000 copies) and the *psen2*^*S4Ter*^/*psen2*^*S4Ter*^ fish (∼1,900 copies) were upregulated. This up-regulation in the *psen2*^*S4Ter*^/*psen2*^*S4Ter*^ fish was clearly significant (p=0.0049), while that in the *psen2*^*S4Ter*^/+ fish was apparent, but not regarded as statistically significant (p=0.0837). The dqPCR raw data is given in supplementary data file S1 Table 1 & 2.

### No increase in DoLA neuron number in embryos homozygous for *S4Ter*

Currently, we do not have an antibody against zebrafish Psen2 protein that would allow us to demonstrate loss of Psen2 activity in homozygous mutants. Also concerning is that a frameshift allele of *psen2* that we have isolated separately, *N140fs*, shows loss of surface melanotic pigmentation when homozygous, suggesting loss of γ-secretase activity (48) whereas homozygous *S4Ter* mutants do not. Therefore, we sought an alternative method to demonstrate loss of *psen2* function due to the *S4Ter* mutation.

Inhibition of *psen2* mRNA translation has been shown to increase the number of a particular spinal cord interneuron – the Dorsal Longitudinal Ascending (DoLA) neuron in zebrafish embryos at 24 hours post fertilization (hpf) (41). Therefore, if *S4Ter* decreases *psen2* function, it might be expected to increase DoLA number (although, as an endogenous mutation rather than blockage of gene expression using a morpholino, *S4Ter* might induce genetic compensation to suppress this phenotype (49)). To examine the effect of *S4Ter* on DoLA number, we collected embryos from a pair-mating of two *psen2*^*S4Ter*^/+ fish to generate a family of embryos comprised, theoretically, of 50% heterozygous mutants, 25% homozygous mutants and 25% wild type genotypes. The embryos were allowed to develop to the 24 hpf stage before *in situ* transcript hybridization against transcripts of the gene *tbx16* that labels DoLA neurons (50). After the number of DoLA neurons in each embryo had been recorded, each embryo was genotyped using PCRs specific for the mutant and wild type alleles. Two-tailed t-tests found no significant differences in DoLA number between any two genotypes. This does not support that *S4Ter* reduces *psen2* activity (Figure 3). Nevertheless, transcriptome analysis (below) shows distinct differences between the brain transcriptomes of *S4Ter* mutant and wild type siblings.

**Figure 3.**
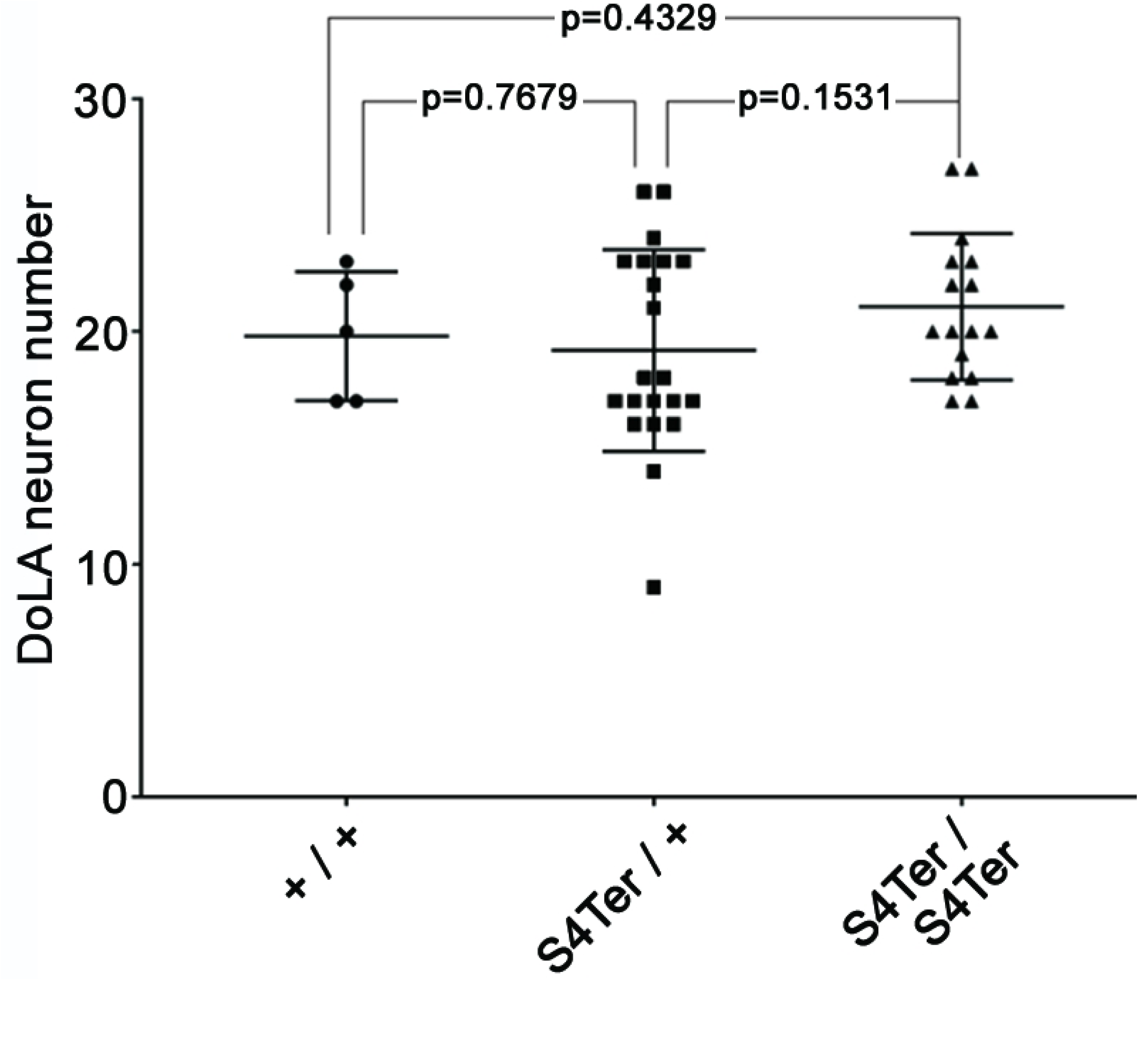
DoLA neuron number assessment of *psen2* activity. 42 embryos at 24 hpf from a pair-mating of a *psen2*^*S4Ter*^/+ female and a *psen2*^*S4Ter*^/+ male were subjected to *in situ* hybridization to detect DoLA neurons that were then counted. Subsequent genotyping of individual embryos revealed 21 heterozygous mutants, 16 homozygous mutants and 5 wild type embryos. Values of p were determined in two-tailed t-tests. Raw data is given in supplementary data file S1 Table 3.

### No apparent effects of the *S4Ter* mutation on transcript splicing

The retention of melanotic pigmentation and normal DoLA neuron number in S4Ter homozygous fish implied that this mutant allele still produces a functional protein. To understand how this might occur, we first checked to see whether the mutation influenced transcript splicing. The *psen2* translation initiation codon and the S4Ter mutation both exist in the second exon of the gene. To test for changes in transcript splicing, we generated cDNA using total RNA purified from wild type and S4Ter homozygous embryos at 24 hours post fertilization (hpf). We then performed PCR using primer pairs binding to sequences in exons 1 and 3, 3 and 7, and 1 and 7 (See Materials and Methods and supplementary data file S1 Appendix 2). Each PCR amplified a single cDNA fragment of the expected size that was essentially identical between the wild type and mutant larvae. (Size differences due to the deletion of 8 nucleotides in the S4Ter mutant allele could not be resolved). Therefore, there is no evidence that the S4Ter mutation changes the splicing of nascent *psen2* transcripts.

### The *S4Ter* mutation allows downstream Met codons to initiate translation

The *S4Ter* mutation might still allow production of a functional protein if a downstream Met codon could act to initiate translation, and the resultant protein retained γ-secretase catalytic activity. Three Met codons exist in the N-terminal-encoding region of the *psen1* ORF (Figure 4A); codons 34, 88, and 97. Codon 34 exists in the cytosolic-coding region before the first transmembrane domain (TM1), while codon 88 codes for a Met residue near the cytosolic surface of TM1 and codon 97’s Met residue is deep within TM1. Since the transmembrane domains of PRESENILIN proteins show high sequence conservation during evolution, we assumed that a protein lacking TM1 could not function so that a functional Psen2 protein might only form if codon 34 or 88 (and remotely possibly 97) were used to initiate translation. Therefore, we fused the known 5’UTR sequences and the first 113 codons of the *psen2* ORF (that includes codons for all TM1 residues) to sequence coding for enhanced green fluorescent protein (EGFP, excluding the EGFP start codon) and incorporated this into the vector pcGlobin2 for synthesis of mRNA. Both wild type and *S4Ter*-mutant versions of this construct were produced (see Figure 4A). Synthetic mRNAs from these vectors were injected into zebrafish embryos at the 1-cell stage and western immunoblots of 24 hpf embryos were subsequently probed with an antibody detecting GFP. This revealed the expected translation initiation from the wild type sequence predominantly at codon 1 (Figure 4B). However, in the presence of the *S4Ter* mutation, there was apparent translation initiation at codon 34 and at either one or both of codons 88 and 97 (although the western immunoblot lacked the resolution to distinguish which, Figure 4B). Therefore, it is highly likely that the *S4Ter* mutant allele produces one or more forms of N-terminally truncated Psen2 protein. Interestingly, these would obey the reading frame preservation rule of the *PRESENILIN* EOfAD mutations and so might reasonably be expected to produce dominant, EOfAD-like effects on brain transcriptomes.

**Figure 4.**
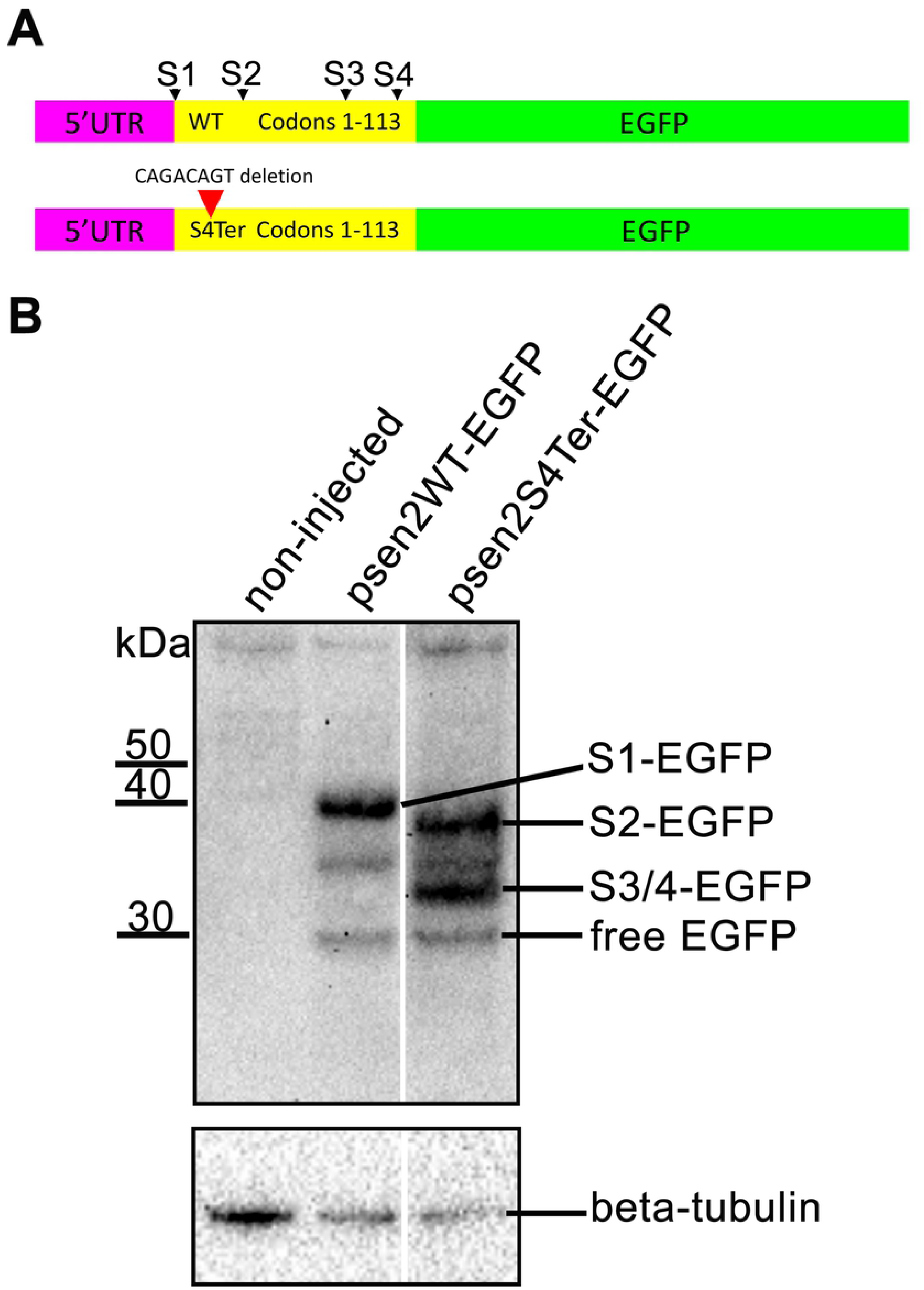
Testing for translation initiation at novel downstream start codons. **A.** The constructs used to test for translation initiation at Met codons downstream of the S4Ter mutation. S1 represents the wild type translation start site and S2-4 are potential downstream translation initiation sites within the first 113 codons. **B.** Western immunoblotting of lysates from embryos injected with the constructs described in A. Translation initiation at S1 (S1-EGFP) does not permit initiation at S2-4. However, in the presence of the S4Ter mutation, initiation is evident at S2 (S2-EGFP) and either S3 or S4 or both (S3/4-EGFP). The identity of the low intensity ∼35kDa protein band is not known but it may be a degradation product of larger fusion protein species. Free EGFP can be produced from EGFP fusion protein species in the lysosome (51, 52).

### Large zebrafish families facilitate reduction of genetic and environmental noise

An advantage of genetic analysis in zebrafish is the ability to reduce genetic and environmental variation in statistical analyses through breeding of large families of siblings that are then raised under near identical environmental conditions (i.e. in the same fish tank or recirculated water aquarium system). The initial heterozygous individual fish identified as carrying *psen2*^*S4Ter*^ was outbred to a wild type fish of the same strain (Tübingen) and then two heterozygous individuals were mated to produce a large family of siblings with wild type (+/+), heterozygous (*psen2*^*S4Ter*^/+), or homozygous (*psen2*^*S4Ter*^/ *psen2*^*S4Ter*^) genotypes. (We have subsequently established a line of fish homozygous for the *psen2*^*S4Ter*^ mutant allele, demonstrating that these fish are both viable and fertile.)

Laboratory zebrafish become sexually mature at between 3 and 5 months of age. Therefore, to examine the transcriptome of young adult zebrafish brains we identified individuals of the desired genotype using PCRs specific for the mutant and wild type alleles on DNA from tail biopsies (“tail clips”) and then removed brains from fish of the desired genotypes at 6 months of age. Total RNA was then purified from these and subjected to either RNA-seq analysis (described below) or digital quantitative PCR (dqPCR) as shown previously in Figure 2).

### RNA-seq data and analysis of *psen2*^*S4Ter*^ effects

To analyze and compare the brain transcriptomes of 6-month-old wild type, heterozygous and homozygous mutant siblings, four female fish of each genotype were examined (Figure 5A). An exploratory principal component analysis (PCA) of gene expression across all samples was generated using gene-level, log_2_-transformed counts per million (Figure 5B), indicating that the difference between wild type (+/+) samples and mutant samples was the dominant source of variability, with PC1 (30.9% of variance) clearly separating the homozygous (Hom) and heterozygous (Het) mutant fish brains from those of the wild type (WT) fish. The Hom and Het mutant fish were largely overlapping with respect to both PC1 and PC2, with the latter accounting for 15.2% of variance in the total dataset.

**Figure 5.**
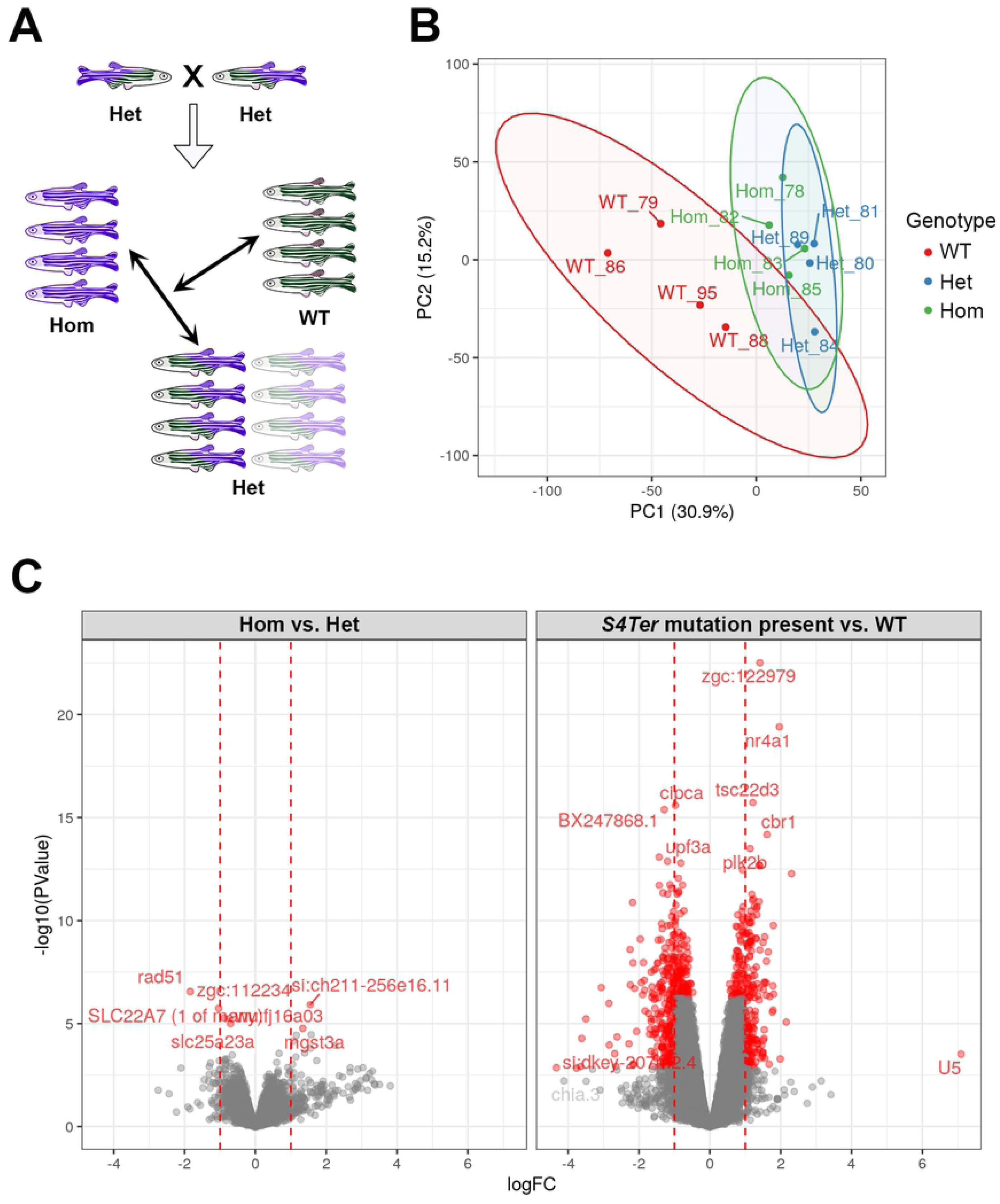
Brain transcriptome analysis. **A**. Pair-mating of two Het zebrafish produces a family made up of WT, Het, and Hom siblings in a ∼1:2:1 ratio respectively. At six months of age, the transcriptomes of entire brains from four female sibling fish each of WT, Het, and Hom genotypes were analyzed. **B.** PCA analysis showing PC1 and PC2 using logCPM values from each sample. The largest source of variability within this dataset was clearly the difference between wild type samples and those containing one or two copies of the *psen2*^*S4Ter*^ allele. **C**. Volcano plots displaying gene differential expression (DE) p-values versus fold-change. Left: DE genes from comparison of Het and Hom brains. Right: DE genes from comparison of brains of fish possessing the *S4Ter* allele (i.e. either Het or Hom) versus WT. The dominant nature of the *S4Ter* mutation is indicated by the relatively restricted differences between Het and Hom transcriptomes compared to *S4Ter* vs. WT.

### Differentially expressed genes (DE genes)

As the Heterozygous and Homozygous *psen2*^*S4Te*^ brains showed very similar patterns of gene expression, we fitted a statistical model for the presence of the mutant allele with an additional term to capture the difference between the two mutant genotypes (Figure 5). In the analysis based on the presence of the mutant allele, genes were considered to be DE using either 1) A Bonferroni-adjusted p-value < 0.01 or, 2) An FDR-adjusted p-value < 0.01 along with an estimated logFC outside of the range ±1. However, as far fewer DE genes were detected when comparing brain transcriptomes between mutant genotypes, a simple FDR of 0.05 was chosen as the DE criterion for that comparison (see supplementary data file S2 Sheet 1 & 3). Ultimately, 615 genes were declared to be DE due to any presence of the *psen2*^*S4Ter*^ mutant allele, while 7 genes were declared to be DE between brains with Hom and Het genotypes. Heatmaps (Figure 6) for the most highly ranked (by FDR) of these DE genes show that the DE genes cluster according to genotype as expected.

**Figure 6.**
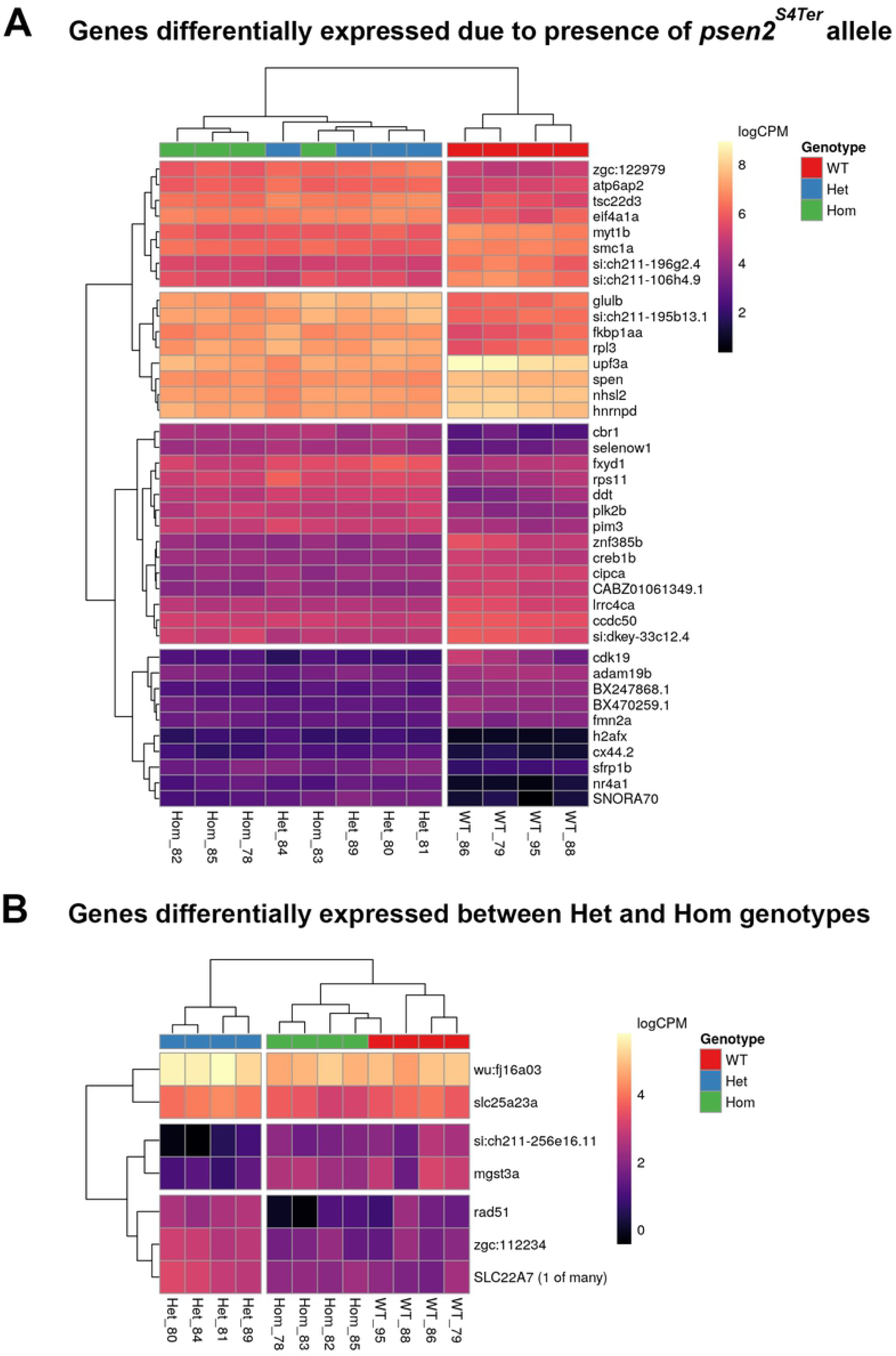
Heatmaps of DE genes. Plotted values are logCPM based on CQN-normalized counts. **A**. The most highly ranked 40 (of the 615) DE genes by FDR identified when comparing WT fish to genotypes possessing the *psen2*^*S4Ter*^ allele. **B**. The 7 most highly-ranked genes (FDR < 0.05) which were detected as DE between Het and Hom mutant genotypes. Unique sample name identifiers are given beneath each column.

### Results from Enrichment Analyses

To predict what changes in molecular/cellular processes might be reflected by the differential expression of genes, enrichment analyses were conducted using the Hallmark, KEGG and GO gene sets defined in the MSigDB database (38). A detailed description of the analyses is publicly available at: https://uofabioinformaticshub.github.io/20170327_Psen2S4Ter_RNASeq/ Full results of enrichment analyses are also listed in supplementary data file S2 sheet 2 & 4. Overall, two strategies for enrichment analysis were used: 1) Testing for enrichment within *discrete sets of DE genes* (WT versus presence of *psen2*^*S4Ter*^ only). 2) Testing for enrichment within the *complete gene lists*, i.e. regardless of DE status, but using direction of fold-change and p-values as a ranking statistic.).

During analysis, we found that the RNA-seq data indicated considerable (and variable) ribosomal RNA (rRNA) content, presumably due to somewhat inefficient depletion of zebrafish rRNA from samples, using a human genome-based rRNA depletion kit. While rRNA sequences could be excluded from the subsequent RNA-seq data bioinformatically, the primary enrichment within DE genes indicated pathways such as *KEGG_RIBOSOME* and *CYTOSOLIC_RIBOSOME* raised concerns that the rRNA contamination might somehow have biased the sampling of cellular RNA, although a mechanism for this is not immediately obvious. Our caution was also raised by recent work showing that gene length might bias the detection of DE gene transcripts in RNA-seq datasets (53). In particular, as a set, ribosomal protein genes have short lengths and may be prone to artefactual identification as DE due to this bias (53). However, when examining the complete gene lists using fry, additional terms which were clearly distinct to rRNA became evident, such as *OXIDATIVE PHOSPHORYLATION* (dre00190), *MITOCHONDRIAL ENVELOPE* (GO:0005740) and *ANION TRANSPORT* (GO:0006820) (Figure 7). As these terms shared virtually no genes with ribosomal-related gene sets, this increases our confidence that these represent real biological differences that are affected by the presence of the *psen2*^*S4Te*r^ allele.

**Figure 7.**
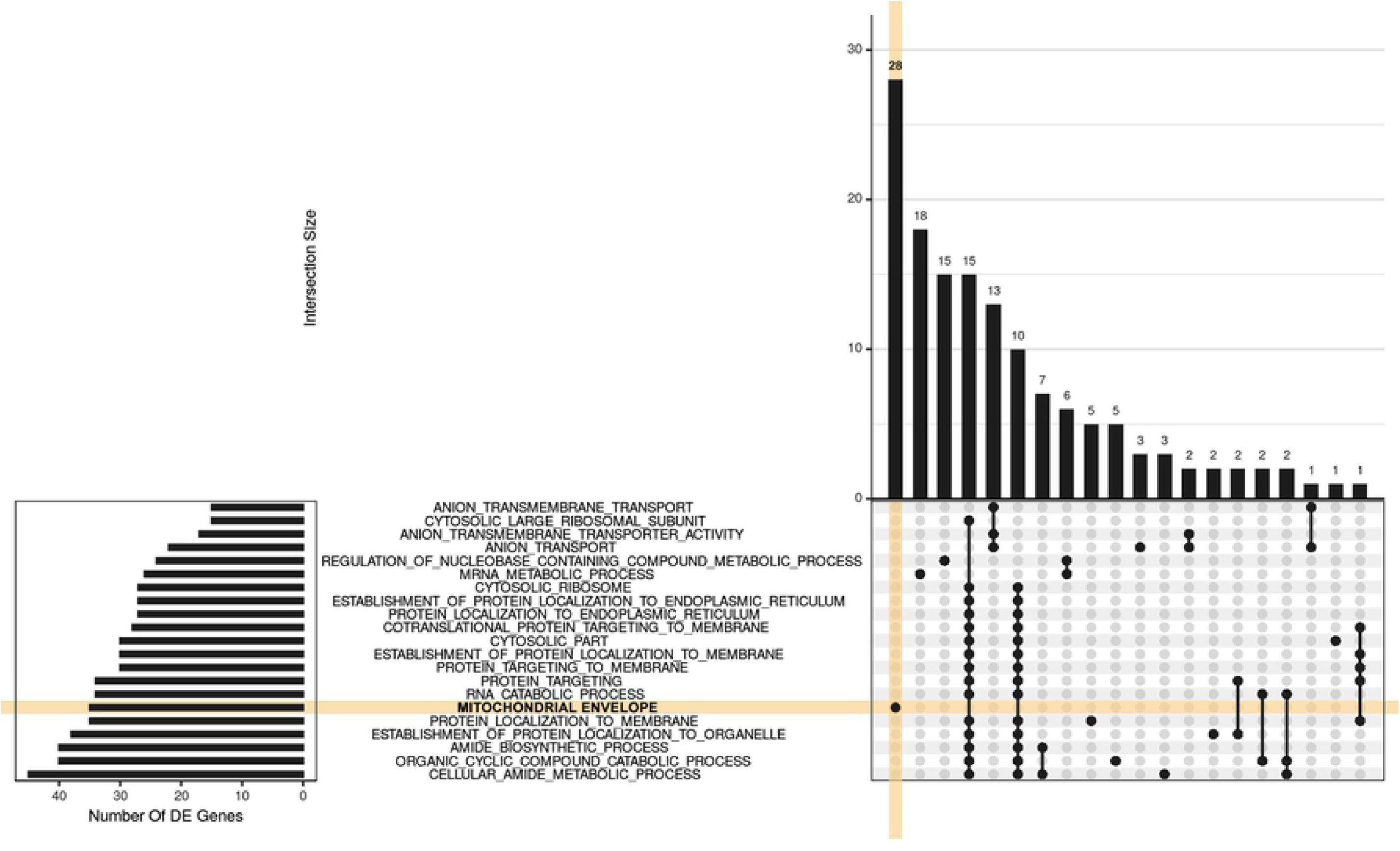
UpSet plot indicating distribution of DE genes within larger significant terms from the GO gene sets. For this visualization, GO terms were restricted to those with 15 or more DE genes, where this represented more than 5% of the gene set, along with an FDR < 0.02 and more than 3 steps back to the ontology root. The 20 largest GO terms satisfying these criteria are shown with the plot being truncated at the right hand side for simplicity. A group of 28 genes is uniquely attributed to the GO *MITOCHONDRIAL ENVELOPE* (orange shading), with a further 18 being relatively unique to the GO *MRNA METABOLIC PROCESS*. The next grouping of 15 genes is unique to GO *REGULATION OF NUCLEOBASE-CONTAINING COMPOUND METABOLIC PROCESS* followed by 25 genes, spread across two clusters of terms which largely represent GO *RIBOSOMAL ACTIVITY*. In between these are 13 genes uniquely associated with the GO *ANION TRANSPORT*.

The presence of the *psen2*^*S4Ter*^ mutation was predicted to affect a number of metabolic pathways, particularly Xenobiotic Metabolism (M5934) from the Hallmark gene sets. However, pathways similar to those seen for an EOfAD-like mutation in the zebrafish *psen1* gene (54), were also observed in the brains of fish with *psen2*^*S4Ter*^, particularly those involving energy production by mitochondria. When homozygous and heterozygous *psen2*^*S4Ter*^ brains were compared, the pathways were even more closely focused around oxidative phosphorylation and mitochondrial function, probably reflecting a critical role for the Psen2 protein in regulating mitochondrial energy production. All pathway enrichment data as generated under fry is given in S2 Sheet 2 & 4.

## Discussion

Well over 200 mutations causing familial Alzheimer’s disease have been identified in the human *PSEN1* and *PSEN2* genes. However, none of these mutations are obviously null (e.g. are frameshift or nonsense mutations that block all mRNAs from producing a protein that includes the normal C-terminal residues) (12). Knowledge of the molecular effects of null mutations in these genes is, therefore, useful since, by exclusion, it could help us determine the functions critically affected by EOfAD mutations.

Alzheimer’s disease takes decades to develop, but we are unable to investigate in detail the molecular changes occurring in the brains of young human carriers of EOfAD mutations since biopsies cannot be taken. Consequently, analysis in animal models is necessary. In this study, we attempted to generate a null mutation in the zebrafish *psen2* gene. We identified an 8-bp deletion in the zebrafish *psen2* gene that forms a premature termination codon (PTC) at the fourth codon position downstream of the start codon (*psen2*^*S4Ter*^). However, this PTC apparently fails to block translation and, instead, reveals cryptic downstream translation start codons that likely drive formation of N-terminally truncated Psen2 protein(s). One or more of these proteins apparently possesses γ-secretase activity since *psen2*^*S4Ter*^ homozygous fish possess melanotic surface pigmentation and normal DoLA neuron numbers in 24 hpf embryos. Translation initiation at cryptic downstream start codons would also explain the lack of NMD of *S4Ter* allele transcripts. The failure of NMD is unlikely to be due to the proximity of the PTC to a downstream exon/exon splice boundary since the nearest such boundary would be far more than 55 nucleotides distant in a spliced mRNA (55).

### The *psen2*^*S4Ter*^ allele shows dominance and may be EOfAD-like

Transcriptome analysis of 6-month-old entire brains from wild type, and *psen2*^*S4Ter*^ heterozygous and homozygous zebrafish showed relatively few differences between the heterozygous and homozygous fish centered around mitochondrial function while extensive changes from wild type were caused by any presence of the *psen2*^*S4Ter*^ allele. The differences that were seen between heterozygous and homozygous brains were in similar functions to those seen for any presence of *psen2*^*S4Ter*^ compared to wild type, only more extreme: effects on mitochondrial function (particularly oxidative phosphorylation) and ribosomal functions. This pattern of effects is most easily explained as *psen2*^*S4Ter*^ displaying a dominant negative phenotype. However, because homozygous fish retain melanotic skin pigmentation, we know that the protein produced by *psen2*^*S4Ter*^ retains some γ-secretase activity (48). Previous research on PRESENILIN protein function suggests two possibilities for the effects of *psen2*^*S4Ter*^ on mitochondria that are not mutually exclusive. In 2009, Area-Gomez et al (23) noted that, in mouse brain, PSEN2 protein is concentrated in the mitochondria-associated membranes of the ER, MAM. In later papers, these researchers noted that loss of *PSEN* gene activity causes increased association of ER membranes with mitochondria (i.e. increased MAM) (56) and decreased oxidative phosphorylation (oxygen consumption) in mouse embryonic fibroblasts (57). While the decreased oxidative phosphorylation could be mimicked by chemical inhibition of γ-secretase, the extent of MAM formation was less sensitive to γ-secretase activity. Increased MAM formation was also observed in fibroblasts from people with EOfAD mutations in *PSEN2*, (or *PSEN1*, or with late onset, sporadic AD) (56). Alternatively, in 2010, Lee et al (58) showed that the Psen1 holoprotein (i.e. not the endoproteolytically-cleaved, γ-secretase-active form of Psen1) was required for normal acidification of lysosomes by promoting N-glycosylation of the V0a1 subunit of v-ATPase that is required for its correct localization to the lysosome. (Note that it has not yet been demonstrated formally that PSEN2 plays a similar role.) Recently, Yambire et al. demonstrated that deficient lysosomal acidification can cause insufficient importation of iron leading to mitochondrial dysfunction (59).

In a 2016 review, we suggested that mutant PRESENILIN holoproteins may interfere with normal holoprotein (putative) multimerization and, thereby, act in a dominant negative manner to interfere with the holoproteins’ normal activity in promoting this N-glycosylation event. (We suggested that the involvement of PRESENILIN holoproteins in multimerization may explain the “reading frame preservation” rule that states that all EOfAD mutations in the *PSEN* genes must preserve an open reading frame that uses the original stop codon (12).) Curiously, both of the PRESENILIN activities mentioned above, the regulation of MAM formation and of endolysosomal acidification, appear to be mediated by the C99 fragment (β-CTF) of APP (57, 60) and this implies that these activities of the PRESENILINs may share a common molecular mechanism.

In conclusion, the *psen2*^*S4Ter*^ mutation is not the null allele we had hoped to isolate and probably results in production of N-terminally truncated Psen2 proteins. These truncated proteins may act in an EOfAD-like manner through their conformity to the “reading frame preservation rule” and display effects on mitochondrial function as we previously observed for an EOfAD-like mutation of *psen1* (54). The *psen2*^*S4Ter*^ mutation also appears to have significant effects on ribosomal functions although an unexpected (and currently inexplicable) correlation of gene differential expression with degree of rRNA contamination in brain RNA samples raises questions around the validity of this result.

## Abbreviations

ACRF: Australian Cancer Research Foundation;
AD: Alzheimer’s Disease;
Cas9: CRISPR associated protein 9;
CPM: counts per million;
CQN: continuous query notification;
CRISPR: clustered regularly interspaced short palindromic repeats;
DE: differentially expressed;
DoLA: dorsal longitudinal ascending neuron;
dNTPs: deoxynucleotide triphosphates;
dqPCR: digital quantitative PCR;
EGFP: enhanced green fluorescent protein;
EOfAD: early onset familial Alzheimer’s disease;
ER: endoplasmic reticulum;
F1/F2/F3: 1^st^/2^nd^/3^rd^ filial generation;
FC: fold change;
FDR: false discovery rate;
GO: Gene Ontology;
GSEA: gene set enrichment analysis;
Hom/HOM: homozygous;
Het/HET: heterozygous;
KEGG: Kyoto Encyclopedia of Genes and Genomes;
MAM: mitochondria-associated membranes (of the ER);
MAPK: MITOGEN-ACTIVATED PROTEIN KINASE;
MD: mean difference;
MsigDB: Molecular Signatures Database;
NHEJ: non-homologous end joining;
NMD: nonsense-mediated mRNA decay;
ORF: open reading frame;
PCR: polymerase chain reaction;
PSEN1: PRESENILIN 1;
PSEN2: PRESENILIN 2;
rRNA: ribosomal RNA;
sgRNA: single guide RNA;
TM1: transmembrane domain 1;
WT: wild type.

## Acknowledgments

The authors wish to thank Seyyed Hani Moussavi Nik for kind assistance in adjusting the conditions for the dqPCR.

## Supporting information

### Supplementary data files

S1 Appendix 1. ***psen2*-EGFP fusion gene sequences**

S1 Table 1. **Expression levels of the wild type *psen2* allele in 25ng total adult brain cDNA**

S1 Table 2. **Expression levels of the *psen2***^***S4Ter***^ **allele in 25ng total adult brain cDNA**

S1 Table 3. **Numbers of DoLA neurons in 24 hpf embryos (revealed by *in situ* transcript hybridization against *tbx16* mRNA)**

S1 Appendix 2. ***psen2***^***S4Ter***^ **transcript splicing tests**

S2 Sheet 1. **DE genes – presence of S4Ter mutation vs. wild type**

S2 Sheet 2. **Pathway enrichment – presence of S4Ter mutation vs. wild type**

S2 Sheet 3. **DE genes – S4Ter Hom vs. S4Ter Het.**

S2. Sheet 4. **Pathway enrichment – S4Ter Hom vs. S4Ter Het.**

